# Light gates hormonal modulation of threat avoidance in female mice

**DOI:** 10.64898/2026.07.08.736856

**Authors:** Marcos L. Aranda, Chase A. Braun, Sophie Hyman, Anika E. Schipma, Tiffany M. Schmidt

## Abstract

Animals must constantly calibrate the costs and benefits of exploration of an environment based on expectations of danger. These decisions are strongly shaped by past experience of perceived threats within that environment and by internal state, which is strongly modulated by circulating gonadal hormones. Although the circuits underlying threat detection are relatively well characterized, how sex hormones shape the long-term behavioral consequences of prior threat experience, and whether this differs across sexes, remains unknown. Here, we show that female mice, like males, exhibit robust long-term threat avoidance (LTTA), avoiding a location where they previously experienced a single visual threat. However, we find that in females this behavior shows strong modulation by the estrous cycle. Surprisingly, we find that though male and female LTTA is driven through glutamate release by the melanopsin-projecting intrinsically photosensitive retinal ganglion cells (ipRGCs) in the thalamic perihabenular nucleus, disruption of this circuit drives completely opposing effects on male versus female LTTA. Moreover, hormonal modulation of LTTA in females requires functional ipRGC input. Thus, despite similar circuit architecture and behavioral outcomes, the individual components of the LTTA circuit play opposing roles in shaping this behavior in males and females, and female LTTA is further tuned by hormonal status.

## Introduction

The calculation of risk and reward in decision making is central to animal survival and reproduction, and thus the circuits underlying these decisions must be optimally calibrated based on past experience, current environmental conditions, and internal state. Visual information is critical in shaping these decisions, because it is involved in not only threat detection, but also in processing contextual cues that allow animals to anticipate and avoid danger in future situations. We have recently defined a new, specialized visual circuit that shapes the degree of threat avoidance based on prior experience involving projections of the melanopsin-expressing, intrinsically photosensitive retinal ganglion cells (ipRGCs) to the perihabenular nucleus (PHb). This long-term threat avoidance (LTTA) circuit is distinct from the well-described threat detection circuits that allow animals to detect and react to threats that appear in their environment and allows animals to use past experience to shape their future decisions around threat avoidance.

Sex differences in threat detection, fear learning, and avoidance behavior are well-documented across species, from invertebrates to humans ^1–6^. Female and male animals often differ not only in the magnitude of their threat responses but also in the behavioral strategies they employ to navigate risky environments. These differences are thought to arise, at least in part, from the actions of gonadal hormones on the neural circuits that translate sensory information and prior experience into adaptive decisions. In female mammals, circulating estrogens and progesterone fluctuate across the reproductive cycle, dynamically modulating synaptic transmission, circuit excitability, and, ultimately, behavior. The risk and reward will be weighted differently based on sexual receptivity, and the optimal balance of these will shift with hormonal cycles. Thus, the capacity to adjust risk-taking versus avoidance behavior accordingly carries a significant evolutionary advantage. However, the circuit-level mechanisms by which gonadal hormones shape these risk/reward calculations and their impact on LTTA is unknown.

In this study, we investigate the interplay between circulating hormones and risk-based decision making in females using our recently-developed LTTA paradigm ^7^, where a single prior exposure to an innately threatening looming stimulus is sufficient to drive selective and long-lasting avoidance of the location where the threat was experienced. Our initial study in males demonstrated that melanopsin signaling is critical for this behavior in males, because males lacking melanopsin fail to avoid the location where the threat was experienced ^7^. Here, we find that while female mice also show LTTA, this behavior is strongly modulated by the presence of circulating hormones. Most surprisingly, however, we find that the loss of melanopsin drives opposing effects on LTTA behavior to that seen in males, despite acting through similar cell types and circuits, and that visual input from ipRGCs is required for hormonal modulation of this behavior.

## Results

### Female mice exhibit measurable long-term threat avoidance (LTTA) behavior

In our recently developed LTTA paradigm we showed that a male mouse will avoid a location where it previously experienced a single exposure to an innately threatening looming stimulus two days prior ^7^. We therefore first wanted to test whether female mice show similar LTTA behavior. Briefly, dark-adapted females were allowed to freely explore an arena beneath an overhead monitor projecting a neutral gray background (see methods). 4 minutes later, mice were exposed to a single 10 second bout of looming stimuli of a given contrast in the center Threat Zone (TZ) of the arena, after which they were placed back in their home cage. 2 days later during the Test phase, animals were assessed for time spent in TZ (relative to Pre-Exposure) while exploring the same arena where they were previously exposed to the looming stimulus, but with no looming stimulus presented (Fig. 1A; see methods).

**Figure 1.**
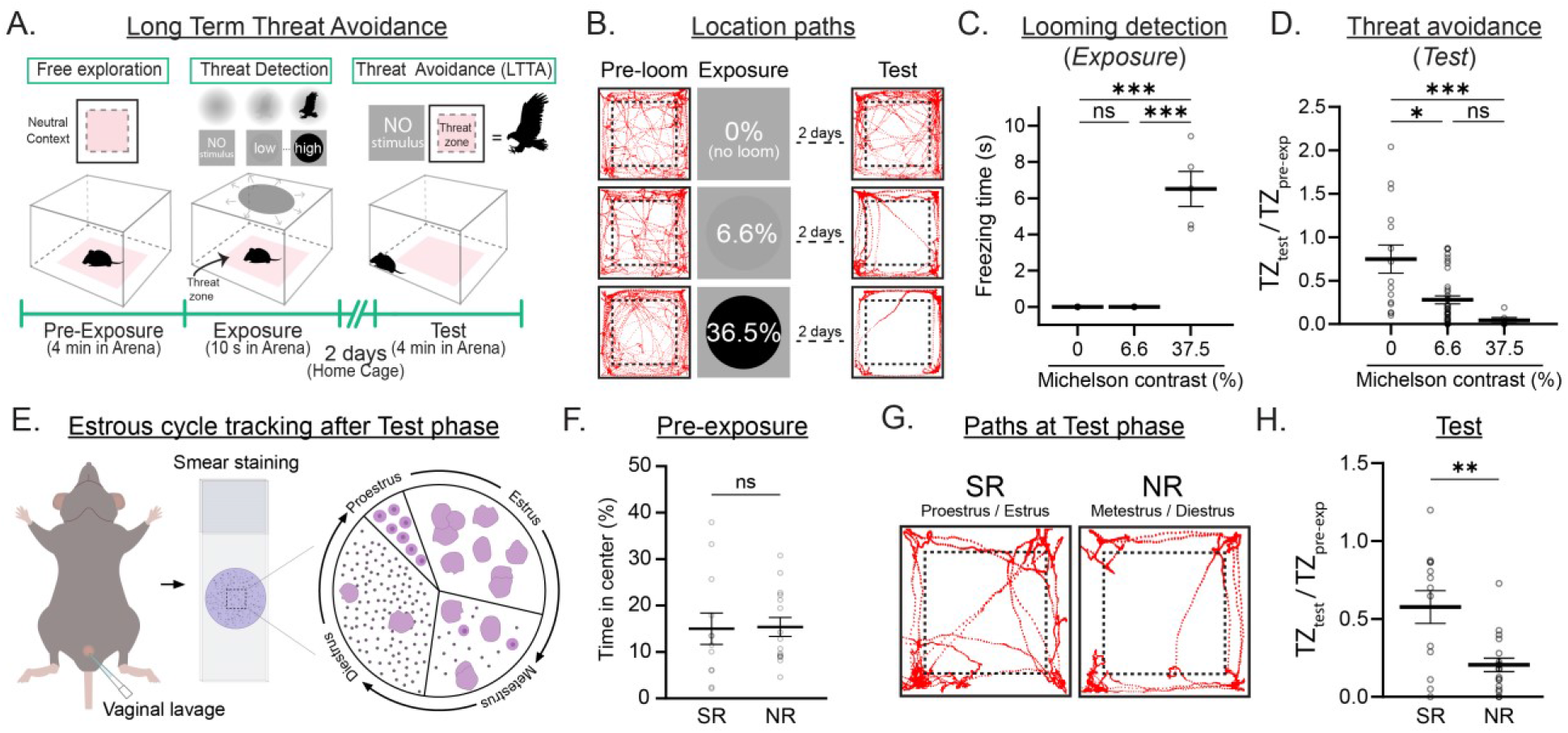
Estrous cycle tunes long-term threat avoidance (LTTA). (A) Schematic representation of LTTA paradigm. (B) Representative location paths of control female mice during the Pre-exposure (Pre-loom) and Test phases of the LTTA paradigm at 0, 6.6 and 36.5% Michelson contrast Exposure. (C) Freezing time at 0% (n = 15 mice), 6.6% (n = 38 mice) and 36.5% (n = 6 mice) Michelson contrast during the Exposure phase of the LTTA paradigm in control female mice (0 vs. 6.6%, P=0.999; 0 vs. 37.5%, P=0.0001; 6.6 vs. 37.5%, P=0.0001). (D) Time in threat zone (TZ) at 0 (n = 15 mice), 6.6 (n = 38 mice) and 36.5% (n = 6 mice) Michelson contrast during the Test phase normalized to time in TZ during the Pre-exposure phase in control male mice (0 vs. 6.6%, P=0.012; 0 vs. 37.5%, P=0.0003; 6.6 vs. 37.5%, P=0.074). (E) Schematic representation of estrous cycle tracking after Test phase of the LTTA paradigm. Proestrus: cells are almost exclusively nucleated epithelial cells. Estrus: cells are predominantly cornified squamous epithelial cells. Metestrus: small darkly stained leukocytes predominate with fragments of cornified squamous epithelial cells and often nucleated epithelial cells. Diestrus: predominating leukocytes and rare epithelial and cornified cells may still be present. Also see Supplementary fig 4. (F) Percentage of time in center during the Pre-exposure phase of WT mice in Sexually Receptive (SR, n = 12) or Non-sexually Receptive (NR, n = 15) phases of estrus (P=0.169). (G) Representative animal position traces of WT female mice in SR and NR estrus stages during the Test phase previously exposed to 6.6% Michelson contrast. (H) Time in TZ during the Test phase normalized to time in TZ during the Pre-exposure phase in SR (n = 12) and NR (n = 15) mice exposed to 6.6% Michelson contrast (P=0.002). All data are Mean ± SEM, n.s. (not significant) P>0.05; *P < 0.05; **P < 0.01; ***P < 0.001. Two-sided Student t- and one-way ANOVA with Kruskal-Wallis’ multiple comparisons test.

We first compared male and female behavior during the Pre-exposure phase and found that male and female mice show similar proportion of exploration of the edges (80% of total time) versus center (20% of total time) of the arena before any threat is presented (Supplementary Fig. 1). We then assessed detection of looming disks varying in contrast from low (6.6% Michelson contrast) to high (36.5%, maximum achievable contrast for a black disk on gray, see methods) by quantifying freezing behavior during the Exposure phase (Fig. 1B-C and Supplementary Fig.1). As in male mice, female mice showed detectable freezing behavior starting at 13.1% Exposure that increased with higher contrast but showed no detectable freezing behavior at 6.6% Exposure (Fig. 1B-C and Supplementary Fig. 1). As in male mice, these results in females defined a subthreshold contrast (6.6%) for freezing behavior and support previous findings that increased looming stimulus contrast enhances saliency to drive increased freezing behavior ^8–10^.

We next quantified long-term avoidance (LTTA) behavior in females by quantifying and comparing time spent in the TZ during the Test versus Pre-exposure phases (Fig. 1A and see methods). Again as in males, female mice previously exposed to a looming stimulus of any contrast, including the lowest level of 6.6%, spent significantly less time in the TZ compared to 0% (i.e. no loom) controls (Fig. 1D, Supplementary Fig. 1 and ^7^). These results demonstrate that female LTTA circuits are more sensitive than the female looming detection circuits and drive long-lasting behavior changes following even a single threat exposure. Importantly, LTTA behavior requires visual input because animals placed in the dark during the Test phase following 6.6% Exposure fail to avoid the TZ, in stark contrast to animals in 10 lux and standard 40 lux illumination (Supplementary Fig. 2). Moreover, female LTTA is not due to increased general anxiety because mice showed no differences 2 days post 0% (no stimulus control) or 6.6% Exposure when assessed in the Elevated Zero Maze (Supplementary Fig. 3). These findings demonstrate that female mice exhibit LTTA behavior and that the circuits underlying LTTA, like those of males, are highly sensitive, long-lasting, and likely distinct from those driving looming-dependent freezing responses and anxiety.

### Estrous cycle tunes LTTA in female mice

Gonadal hormones are powerful modulators of the neural circuits that govern threat processing ^11–14^. In female mice, circulating sex hormones fluctuate across the estrous cycle, peaking during the proestrus/estrus phases, which correspond to the sexually receptive (SR) state, and reaching their nadir during metestrus/diestrus, the non-sexually receptive (NR) state. Thus, hormonal cycles could impact both threat detection and threat avoidance. We first examined whether estrous phase influences looming stimulus detection itself, taking advantage of the Threat Contrast Sensitivity (TCS) paradigm ^9^, which allows us to measure distinct behavioral responses as mice are exposed to looming stimuli of varying contrast (6.6, 10.1, 13.1, 21.9, 27.5, and 36.5% Michelson contrast). We found that SR- and NR-female mice showed no difference in behavior, during either Pre-exposure or looming exposure, to any of the looming contrasts presented, indicating that hormones do not modulate threat responses or general exploratory behavior of the arena (Supplementary Fig. 4).

We next examined whether estrous cycle state impacts LTTA behavior by comparing SR and NR female LTTA following 6.6% Michelson contrast Exposure to isolate impacts on LTTA circuits from those of the less sensitive looming detection circuit (Fig. 1 and ^7^). We found that NR-females spent significantly less time in the TZ compared to SR-females in the Test phase, despite similar behavior during Pre-exposure and Exposure phases (Fig. 1F-H). These results demonstrate that estrus significantly impacts threat avoidance but not baseline exploration or threat detection itself.

If estrous stage modulates LTTA, then removal of circulating hormones should eliminate the differences between SR and NR females. Thus, as a second test of hormonal impact on female LTTA, we assessed whether loss of circulating hormones impacts female LTTA. To do this, we ovariectomized (Ovx) female mice to eliminate the major source of circulating estrogen and progesterone, which mimics mimic NR stages of the estrous cycle. We then exposed Ovx and *Sham* SR- and *Sham* NR-female control littermates to the LTTA paradigm following 6.6% Exposure (Fig. 2A, see methods). All groups showed similar baseline exploratory behavior during Pre-Exposure, supporting our findings above that circulating gonadal hormones are dispensable for this behavior. In agreement with our hypothesis, Ovx females spent significantly less time in the TZ than *Sham-*SR females but were similar to *Sham-*NR females (Fig. 2B-D). These results indicate that ovarian-derived sex hormones are required for estrus-dependent effects on LTTA.

**Figure 2.**
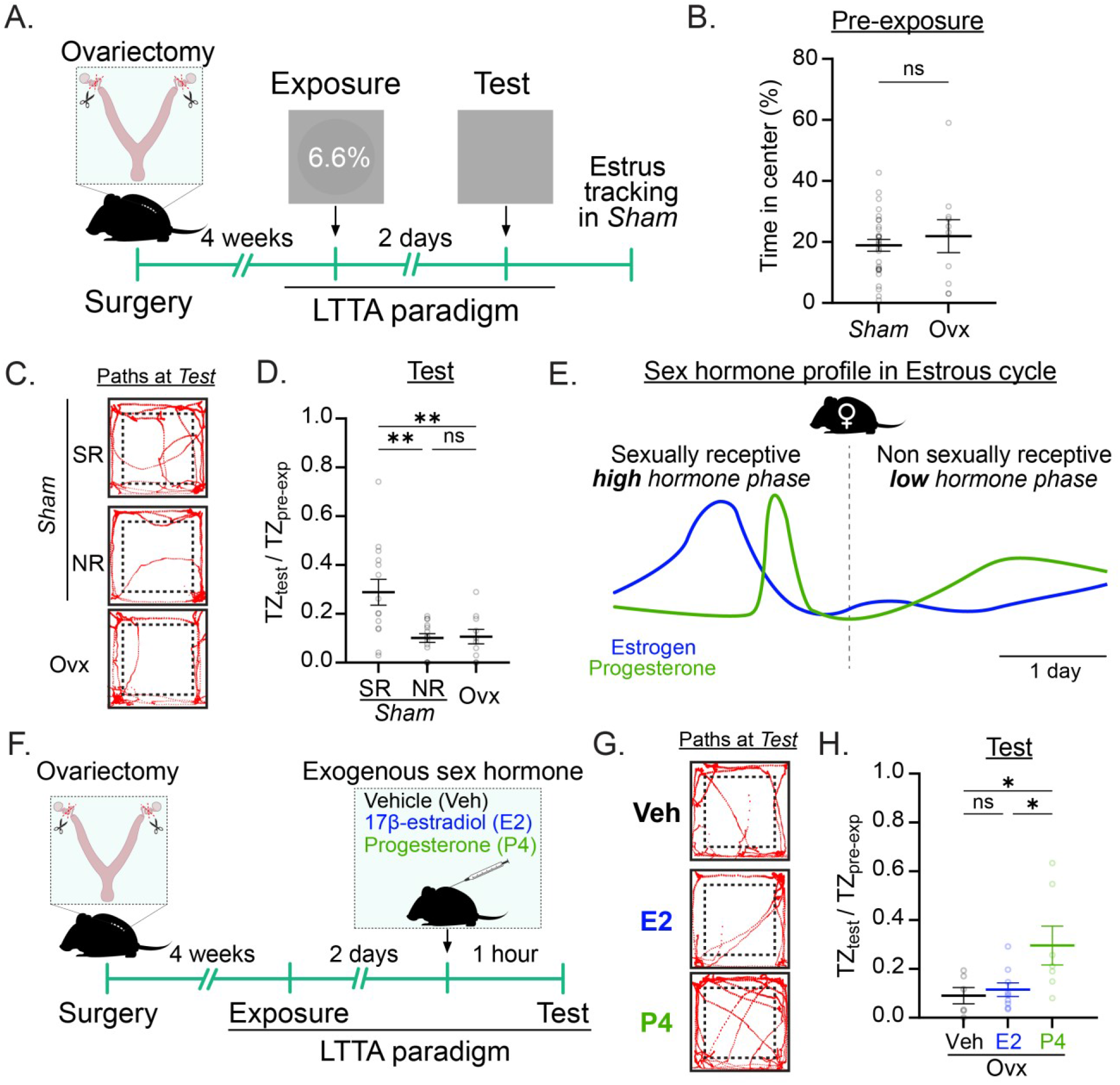
Progesterone drive estrus-biased LTTA. (A) Schematic experimental timeline of ovariectomy followed by the LTTA paradigm. (B) Percentage of time in center during the Pre-exposure phase of *Sham* (n = 28) or Ovariectomized (Ovx, n = 10) WT female mice (P=0.041). (C) Representative animal position traces of *Sham* in Sexually Receptive (SR, n = 14) or Non-sexually Receptive (NR, n = 14) phases of estrus and Ovx (n = 10) females during the Test phase previously exposed to 6.6% Michelson contrast. (D) Time in threat zone (TZ) during the Test phase normalized to time in TZ during the Pre-exposure phase in *Sham* SR (n = 14), *Sham* NR (n = 14) and Ovx (n = 10) female mice exposed to 6.6% Michelson contrast (*Sham* SR vs. *Sham* NR, P=0.002; *Sham* SR vs. Ovx, P=0.007; *Sham* NR vs. Ovx, P=0.994). (E) Schematic representation of estrogen and progesterone levels in SR and NR stages. (F) Schematic experimental timeline of ovariectomy followed by exogenous Vehicle (Veh, corn oil), 17β-estradiol (E2) or Progesterone (P4) subcutaneous administration before the Test phase of the LTTA paradigm in control female mice. (F) Representative animal position traces of Veh, E2 and P4 injected-Ovx females during the Test phase previously exposed to 6.6% Michelson contrast. (H) Time in TZ during the Test phase normalized to time in TZ during the Pre-exposure phase of Veh (n = 6), E2 (n = 9) and P4 (n = 7) control injected-Ovx female mice exposed to 6.6% Michelson contrast (Veh vs. E2, P=0.938; Veh vs. P4, P=0.036; E2 vs. P4, P=0.042). All data are Mean ± SEM, n.s. (not significant) P>0.05; *P < 0.05; **P < 0.01. Two-sided Student t-test and one-way ANOVA with Tukey’s multiple comparisons test.

### Progesterone is required for estrus-dependent tuning of LTTA

We next sought to determine which circulating gonadal hormone(s) modulate LTTA behavior. Among the gonadal hormones that fluctuate across the estrous cycle, estrogens and progesterone are well-established modulators of neural circuit function and behaviors ^11–14^ (Fig. 2E). To define the role of estrogen and progesterone in modulating LTTA, we compared LTTA behavior in Ovx mice where estrogen or progesterone were administered prior to the Test phase of LTTA. Specifically, we subcutaneously (s.c.) administered Vehicle (Veh, corn oil) to all animals 1 hour prior to the Pre-exposure, and Veh, 17β-estradiol (E2) or progesterone (P4) to Ovx control female mice 1 hour prior to the Test phase of the LTTA paradigm (Fig. 2F, see methods). Interestingly, while no differences were observed between Veh- and E2-injected mice, P4 administration resulted in increased time spent in the TZ that matched that of SR-females (Fig. 2G-H). These results suggest that progesterone, but not estrogen, is required for estrus-dependent impacts on female LTTA.

### Melanopsin is required for female LTTA

The melanopsin-(*Opn4*)-expressing intrinsically photosensitive retinal ganglion cells (ipRGCs) (Fig. 3A) tune LTTA in male mice, with melanopsin-null male mice spending significantly more time in the TZ during the Test phase, indicating that melanopsin facilitates TZ avoidance ^7^. Given that male and female mice both show LTTA behavior, we hypothesized that melanopsin signaling would tune female LTTA in a similar manner. To test this, we compared TZ avoidance of melanopsin null (*Opn4^-/-^*) and littermate control females following 6.6% Exposure in the LTTA paradigm. Surprisingly, though male *Opn4^-/-^* animals spend significantly more time in the TZ during the Test phase (Supplementary Fig. 5 and ^7^), female *Opn4^-/-^* animals spent significantly less time in the TZ (Fig. 3B-C), indicating that melanopsin tunes LTTA in the opposite direction in female mice. Importantly, *Opn4^-/-^* and control littermate female mice show similar baseline exploratory behavior during Pre-exposure phase (Supplementary Fig. 5), indicating that differences in exploration do not drive observed differences in LTTA. These results show that melanopsin in intact females serves to dampen LTTA and promote exploration, while in males, melanopsin actually enhances LTTA and drives TZ avoidance, revealing a surprising, and opposing, sex-dependent impact of melanopsin.

**Figure 3.**
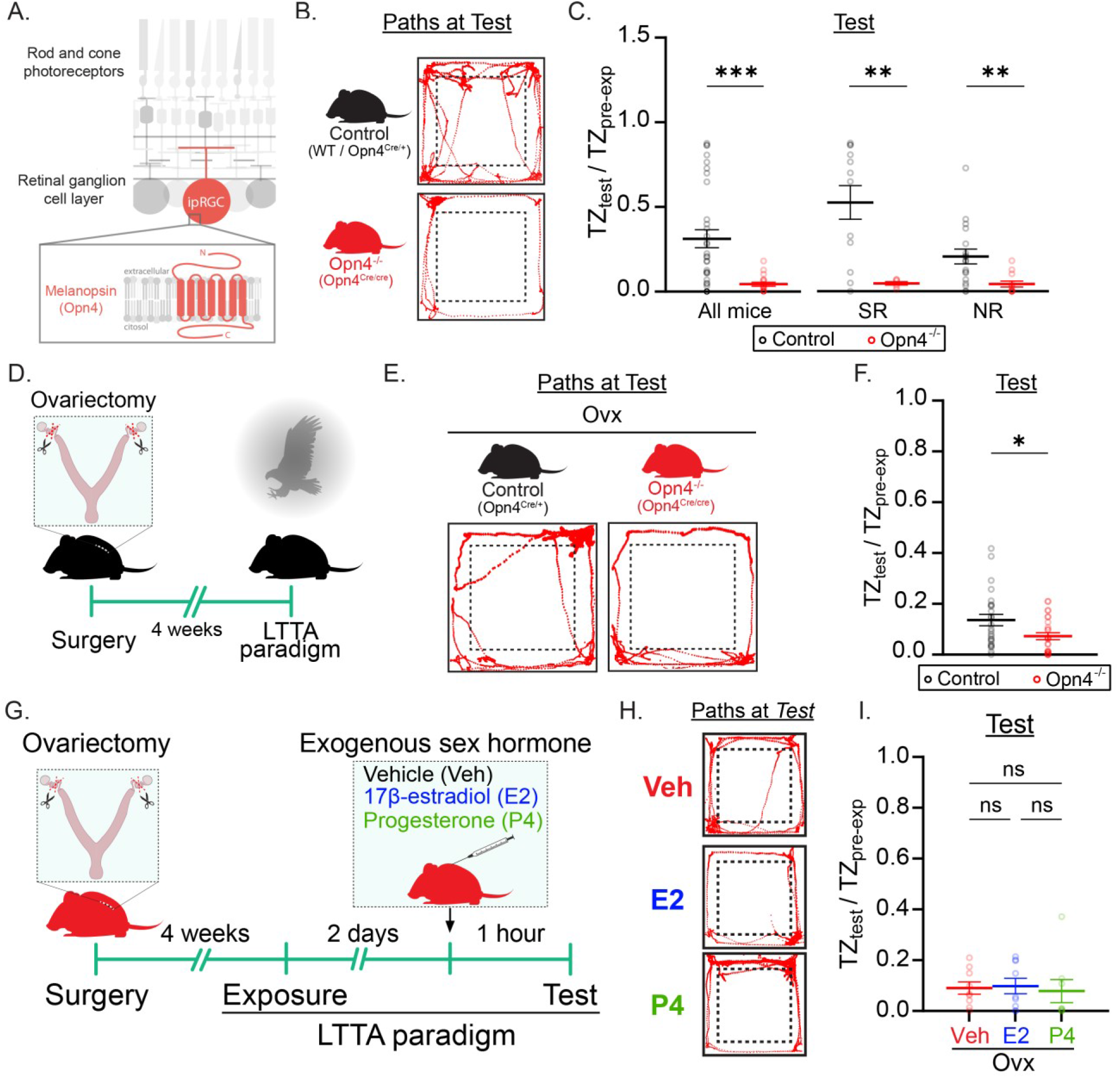
ipRGCs tunes LTTA in female mice. (A) Scheme of melanopsin expression in ipRGCs. (B) Representative animal position traces of control and melanopsin-null (*Opn4^-/-^*) female mice during the Test phase of the LTTA previously exposed to 6.6% Michelson contrast. (C) Time in threat zone (TZ) during the Test phase normalized to time in TZ during the Pre-exposure phase in control (n = 32, black) and *Opn4^-/-^*(n = 18, red) combined (All mice, P=0.0002) or split into Sexually Receptive (SR, Control: n=12, *Opn4^-/-^*: n=6) or Non-sexually Receptive (NR, Control: n=18, *Opn4^-/-^*: n=12) estrous stages exposed to 6.6% control mice Michelson contrast (Control SR vs. *Opn4^-/-^*SR, P= 0097; Control NR vs. *Opn4^-/-^* NR, P= 0015). (D) Schematic representation of ovariectomy followed by the LTTA paradigm in control and *Opn4^-/-^* female mice. (E) Representative animal position traces of control and *Opn4^-/-^* Ovx-female mice during the Test phase previously exposed to 6.6% Michelson contrast. (F) Time in TZ during the Test phase normalized to time in TZ during the Pre-exposure phase of control (n = 25, black) and *Opn4^-/-^* (n = 26, red,) Ovx-female mice (P=0.021). (G) Schematic experimental timeline of ovariectomy followed by exogenous Vehicle (Veh, corn oil), 17β-estradiol (E2) or Progesterone (P4) subcutaneous administration before the Test phase of the LTTA paradigm in *Opn4^-/-^* female mice. (H) Representative animal position traces of Veh, E2 and P4 *Opn4^-/-^* injected-Ovx females during the Test phase previously exposed to 6.6% Michelson contrast. (I) Time in TZ during the Test phase normalized to time in TZ during the Pre-exposure phase of Veh (n = 9), E2 (n = 9) and P4 (n = 8) *Opn4^-/-^* injected-Ovx female mice exposed to 6.6% Michelson contrast (Veh vs. E2, P=0.985; Veh vs. P4, P=0.967; E2 vs. P4, P=0.914). All data are Mean ± SEM, n.s. (not significant) P>0.05; *P < 0.05; **P < 0.01; ***P < 0.001. Two-sided Student t- and Mann-Whitney U tests, and one-way ANOVA with Tukey’s multiple comparisons test.

Glutamate release from ipRGCs is essential for male LTTA, and removal of ipRGC glutamate release in male mice drives a parallel phenotype to that of *Opn4^-/-^* males, leading to increased time in TZ during the Test phase in LTTA ^7^. Given the opposing effects of ipRGCs on male and female LTTA, we next tested whether removal of *Vglut2* specifically from ipRGCs impacts female LTTA. To do this, we compared LTTA in Vglut2cKO (conditional knockout; *Opn4^Cre/+^; Vglut2^fl/fl^*) and control littermate (*Opn4^+/+^; Vglut2^fl/fl^*) females where *Vglut2* is removed specifically from ipRGCs. As with *Opn4^-/-^* females, we found that female Vglut2cKO females showed decreased time in TZ despite normal exploratory behavior during the Pre-exposure phase (Supplementary Fig. 5). Thus, though males and females both exhibit similar LTTA behavior, and though male and female LTTA requires melanopsin signaling and glutamate release from ipRGCs, genetic manipulations reveal fundamental differences in the circuit mechanisms because ipRGC glutamate release modulates male versus female LTTA behavior in opposing directions.

### Melanopsin signaling is required for estrous-dependent impacts on LTTA

We next asked whether melanopsin is required for hormonal impacts on female LTTA. To test this, we compared LTTA of control and *Opn4^-/-^*female littermates split based on SR versus NR stages. Surprisingly, *Opn4^-/-^*females showed similar decreased time in TZ during the Test Phase regardless of estrous stage, indicating that melanopsin may be required for estrus-dependent effects on LTTA.

As a second test of melanopsin’s requirement for gonadal hormone modulation of female LTTA, we next compared LTTA in Ovx-*Opn4^-/-^* versus Ovx-control littermate females (Fig. 3D). Ovx-*Opn4^-/-^* females showed decreased time in TZ compared to Ovx-control littermates during the Test phase (Fig. 3E-F) despite similar exploratory behavior Pre-exposure (Supplementary Fig. 5), indicating that melanopsin dampens LTTA even in the absence of circulating sex hormones.

We next tested whether exogenous addition of progesterone or estrogen could modulate LTTA in *Opn4^-/-^*females. To test this, we compared LTTA in Ovx-*Opn4^-/-^* female littermates where estrogen, progesterone, or vehicle were administered during the Test phase. Specifically, we s.c. administered Veh to all Ovx-*Opn4^-/-^*females 1-hour prior to the Pre-exposure phase, and then either Veh, E2 or P4 1 hour prior to the Test phase (Fig. 3G, see methods). We observed no differences in LTTA behavior between Veh-, E2- or P4-injected Ovx-*Opn4^-/-^*female mice (Fig. 3H-I), indicating that sex hormones do not alter LTTA behavior in the absence of melanopsin signaling. Collectively, these results show that melanopsin signaling in females is required for hormonal modulation of LTTA behavior. Consistent with this, progesterone receptors (*Pgr*) are expressed in ipRGCs, while estrogen receptors (*Esr1*, *Esr2* and *Gper1*) show comparatively low expression in this cell population (Supplementary Fig. 6 and ^15^), suggesting that ipRGCs may represent one direct molecular target through which progesterone could modulate LTTA.

### The perihabenular nucleus is required for tuning LTTA in female mice

In our recent study in males, we found that the ipRGC-perihabenular nucleus (PHb) circuit is required for turning LTTA in male mice ^7^, defining distinct circuits for threat avoidance and threat detection. Given the differing phenotypes in male and female *Opn4^-/-^* and Vglut2cKO animals, we next examined whether ipRGCs impact male and female LTTA through an ipRGC to PHb circuit. ipRGCs primarily innervate GABAergic neurons in the caudal PHb (PHb^GABA^), which in turn make local connections with excitatory-relay and anterior PHb^GABA^ neurons ^16–18^. If PHb^GABA^ neurons are required for LTTA, then silencing that specific population should alter LTTA in female mice. To test this, we injected Cre-dependent Gi-DREADDs into the PHb of *Gad2^Cre^* female mice to silence PHb^GABA^ (Fig. 4A-B, see Supplementary Table 1) neurons. We then intraperitoneally (i.p.) injected either vehicle or the DREADD agonist clozapine-N-oxide (CNO) 30 minutes prior to both the Pre-exposure and Test phases of the LTTA paradigm (Fig. 4C). Silencing of PHb^GABA^ (Fig. 4D-F) neurons during the Test phase increased LTTA, causing animals to spend significantly less time in the TZ, confirming that the GABAergic neurons of the ipRGC-recipient PHb are required for LTTA. Importantly, i.p. CNO injection did not alter exploratory behavior during Pre-exposure phase compared to vehicle-injected mice, indicating that i.p. CNO itself has no effect (Fig. 4D). Notably, this decreased time in TZ following silencing of PHb^GABA^ neurons is completely the opposite of that previously reported for males ^7^, indicating that though the same basic ipRGC-PHb circuit tunes male and female LTTA, it does so with opposing impacts on TZ avoidance.

**Figure 4.**
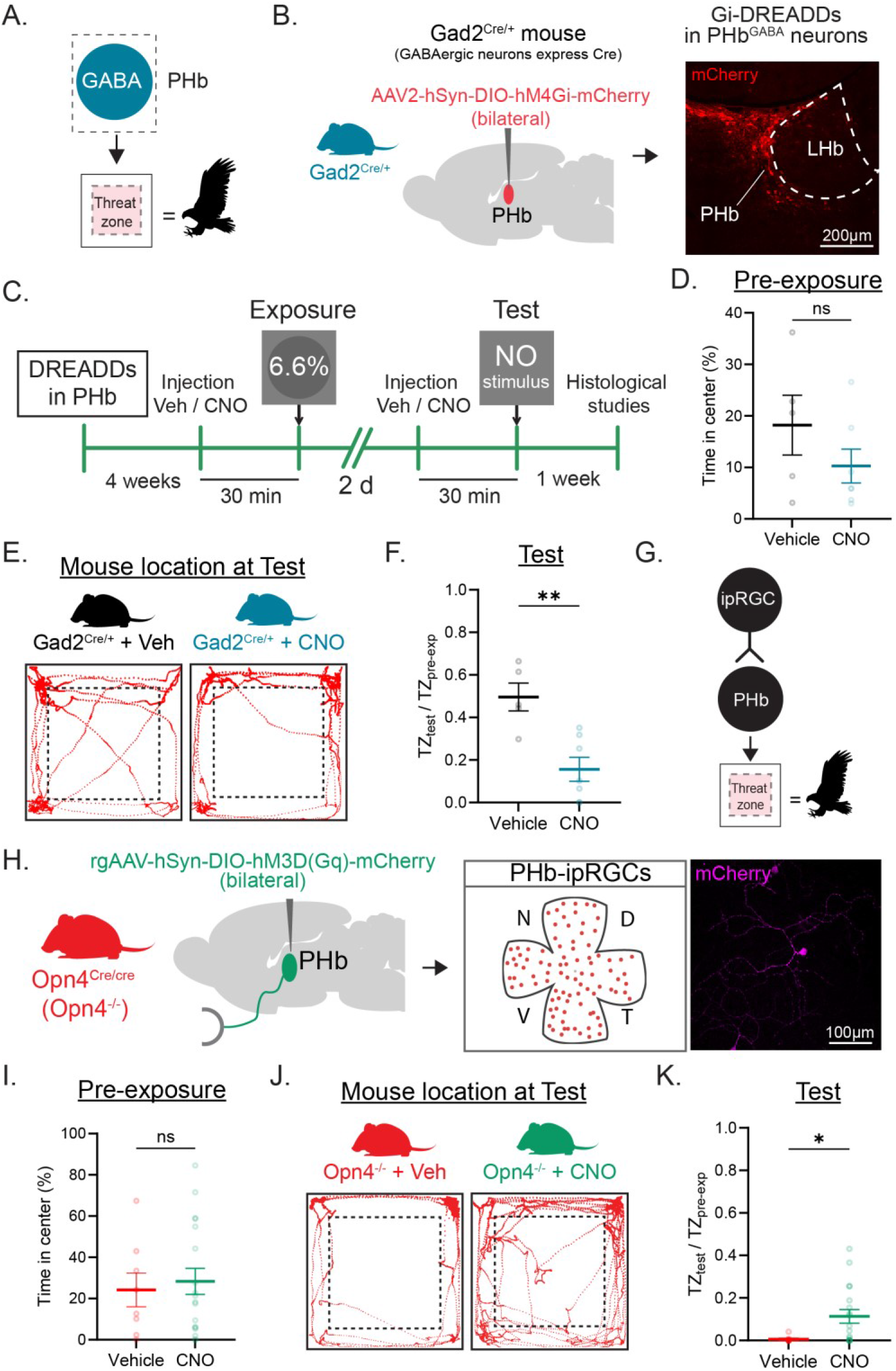
ipRGC-PHb^GABA^ circuit is required for LTTA. (A) Schematic representation of PHb^GABA^ neurons driving LTTA. (B) Scheme of viral injection of Gi-DREADDs in PHb of *Gad2^Cre^* female mice (left) and representative image of Gi-DREADDs expression in PHb (right). (C) Schematic experimental timeline for DREADDs injections in PHb in female mice. (D) Percentage of time in center during the Pre-exposure phase of Vehicle (n = 5, black) or CNO-injected (n = 7, blue) *Gad2^Cre^* female mice (P=0.321). (E) Representative *Gad2^Cre^* animal position traces during the Test phase of the LTTA paradigm after 6.6% exposure Michelson contrast. (F) Time in the threat zone (TZ) during the Test phase normalized to time in TZ during the Pre-exposure phase of Vehicle (n = 5, black) or CNO-injected *Gad2^Cre^* (n = 7, blue) female mice (P=0.003). (G) Schematic representation of ipRGC-PHb circuit driving LTTA^7^. (H) Scheme of viral injection of rgGq-DREADDs in PHb of *Opn4^-/-^* female mice (left) and representative images of Gq-DREADDs expression in a PHb-ipRGC (right). (I) Percentage of time in center during the Pre-exposure phase of Vehicle- (n = 9, red) or CNO-injected (n = 18, green) *Opn4^-/-^*female mice (P=0.859). (J) Representative *Opn4^-/-^* animal position traces during the Test phase of the LTTA paradigm after 6.6% exposure Michelson contrast. (K) Time in TZ during the Test phase normalized to time in TZ during the Pre-exposure phase of Vehicle (n = 9, red) or CNO-injected *Opn4^-/-^*(n = 18, green) female mice (P=0.019). All data are Mean ± SEM, n.s. (not significant) P>0.05; *P < 0.05; *P < 0.01. Two-sided Student t- and Mann-Whitney U tests.

Light activates melanopsin in ipRGCs, triggering a Gq-mediated phototransduction cascade that ultimately depolarizes the cells and drives glutamate release onto downstream targets^19^. If PHb-ipRGCs are required for LTTA, then activation of PHb-ipRGCs in *Opn4^-/-^* females should rescue the observed decreased time in TZ. We have previously shown that activation of Gq-DREADDs in ipRGCs restores melanopsin-like photoresponses in *Opn4^-/-^* ipRGCs *ex vivo* ^20^ and rescues *in vivo* ipRGC-dependent behavior in the LTTA paradigm ^7^. We therefore sought to rescue female *Opn4^-/-^*LTTA behavior using this approach. Specifically, we bilaterally injected retrograde (rg)AAV into the PHb of melanopsin null (*Opn4^Cre/Cre^*) female animals to drive Cre-dependent expression of Gq-DREADDs (rgAAV-hSyn-DIO-hM3Gq-mCherry) selectively in PHb-ipRGCs (Fig. 4G-H, see Supplementary Table 1). We then intraperitoneally injected either vehicle or CNO 30 minutes prior to both the Pre-exposure and Test phases (Fig. 4C). Consistent with our prediction, CNO-injected mice showed decreased LTTA, spending significantly more time in the TZ than vehicle-injected *Opn4^-/-^* female littermates (Fig. 4I-K). Importantly, CNO- versus vehicle-injected *Opn4^-/-^* females showed no differences in baseline exploratory behavior (Fig. 4I), again indicating that CNO itself does not alter exploratory behavior. These data indicate that Gq-DREADD activation of PHb-ipRGCs restores baseline LTTA in *Opn4^-/-^* female mice, demonstrating that PHb-ipRGCs are required for female LTTA. Notably, this ability to rescue the behavioral phenotype also suggests that *Opn4^-/^* behavioral changes are not due to extensive developmental rewiring because the behavior can be restored acutely and in adulthood.

### Estrous cycle tunes PHb neuron activity during LTTA

Our previous study showed that the ipRGC to PHb signaling is required to facilitate the recall of the threat/context association in the LTTA paradigm ^7^. To understand how estrous stages impact PHb signaling dynamics during LTTA, we used fiber photometry (FP) to measure *in vivo* bulk Ca^2+^ activity in the PHb in estrus-tracked female littermates during the Test phase of the LTTA paradigm. To do this, we injected an AAV containing GCamp6s (AAV1-CAG-GCaMP6s-WPRE-SV40) and implanted a FP canula unilaterally in the PHb of female mice (Fig. 5A-B, see Supplementary Table 1 and methods). We analyzed the activity in the PHb while mice occasionally transitioned from the edge to the TZ and vice versa during the Test phase (Fig. 5C). We found a significant decrease in the calcium signal z-score in the PHb when NR-female mice transitioned from the edge to TZ, but no significant change in SR-females (Fig. 5 D-F), indicating the PHb neuron activity signals entry into the TZ only during NR stages of estrus where circulating sex hormone levels are low. Interestingly, no change was observed when either NR or SR-females transitioned from the TZ to the edges (Fig. 5G-I), suggesting that PHb neurons signal entry into the area where the threat was experienced.

**Figure 5.**
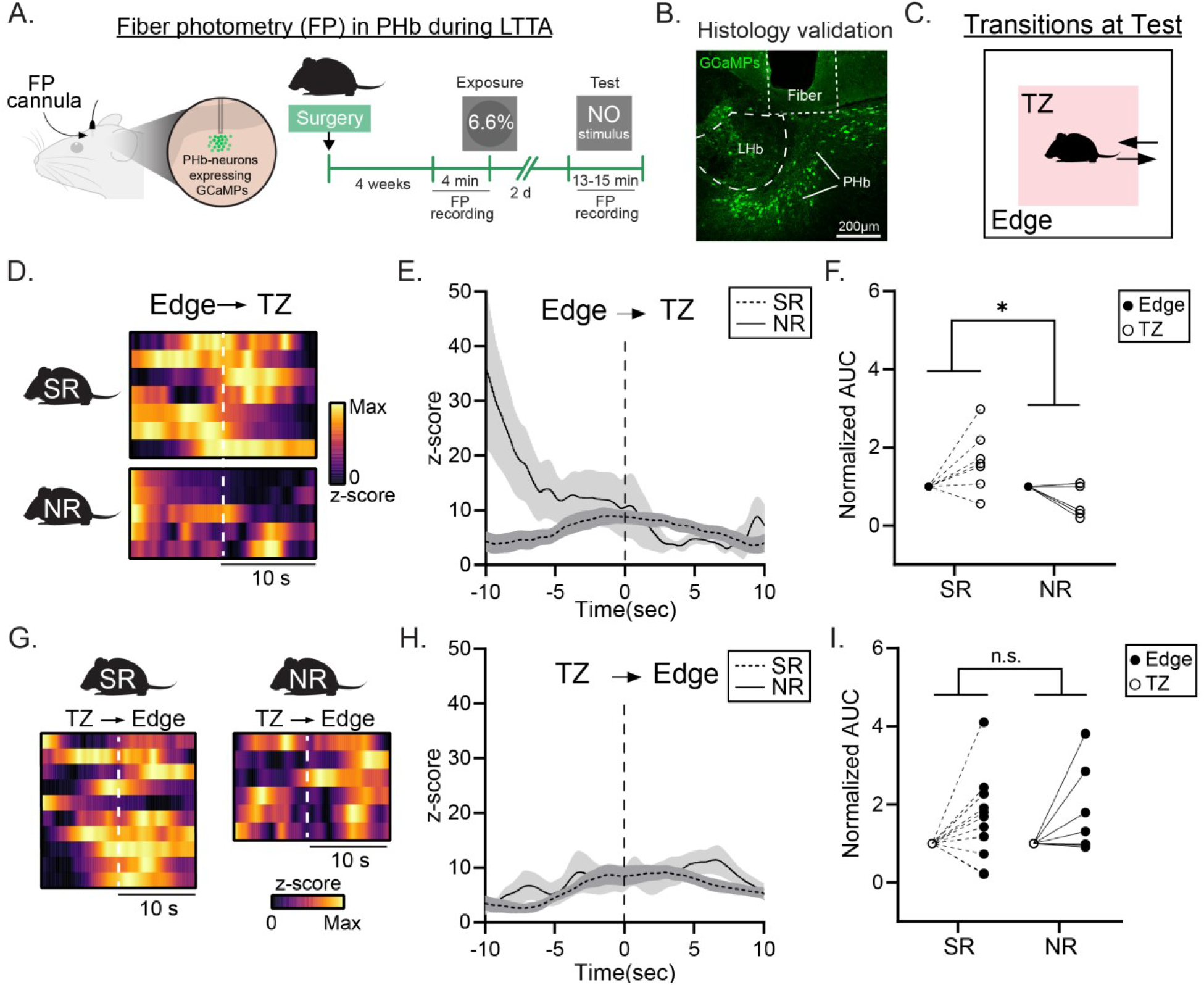
Estrus stages tune PHb neuron activity during LTTA. (A) Scheme of experimental design for fiber photometry (FP) recordings of control female mice. (B) Representative image of GCaMPs expression and optic fiber implantation in the PHb. (C) Schematic representation of transitions from edge to the threat zone (TZ) and TZ to edge during the Test phase of the LTTA paradigm. (D-F) heatmaps (D), *z*-score of FP signal (E) and AUC of calcium signal z-score (F) when mice transition from edges to TZ during the Test phase of the LTTA paradigm in SR (n = 7, dashed line) and NR (n = 5, continuous line) control female mice (P=0.046). (G-J) heatmaps (G), *z*-score of FP signal (H) and AUC of calcium signal z-score (I) when mice transition from TZ to edge during the Test phase of the LTTA paradigm in SR (n = 10, dashed line) and NR (n = 6, continuous line) control female mice (P=0.797). All data are Mean ± SEM, n.s. (not significant) P>0.05; *P < 0.05. Two-sided paired Student t-tests.

## Discussion

The ability to balance risk-taking against safety is fundamental to animal survival, and how this balance is tuned by sex and reproductive state remains poorly understood. Here, we demonstrate that female mice exhibit robust long-term threat avoidance behavior in the LTTA paradigm, and will avoid the TZ for days after a single exposure to an innately threatening visual looming stimulus. Importantly, as in male mice, this behavior in females is distinct from canonical looming-evoked freezing and anxiety circuits, and similarly requires melanopsin-expressing ipRGC input to the PHb nucleus. Together, these findings establish that LTTA is conserved across sexes while revealing important sex-specific tuning of the underlying circuit.

Both male and female LTTA requires melanopsin signaling and glutamate release from ipRGCs as well as PHb^GABA^ neurons, indicating similar cell types and circuitry are involved in the behavior. However, our studies in male and female mice unveil a marked sex difference in the underlying circuit mechanisms of LTTA behavior. When the ipRGC-PHb circuit is dysfunctional in males, animals show decreased LTTA, spending more time in the TZ ^7^. Interestingly, disrupting the same circuit in females, whether through *Opn4^-/-^*, Vglut2cKO, or chemogenetic manipulations, drives increased LTTA, causing animals to spend even less time in the TZ. Notably, this opposing directionality is not apparent at the behavioral level: male and female mice show broadly similar LTTA. However, these sex-dependent differences are unmasked upon circuit manipulation, suggesting that the ipRGC-PHb circuit is engaged differently across sexes despite producing a convergent behavioral output. These findings highlight the importance of studying circuit mechanisms in males and females because fundamentally different circuit mechanisms can lead to outwardly similar behaviors. Notably, circuit differences in males and females are not due to hormonal modulation, because differences in ipRGC influence persist in the absence of circulating sex hormones. Identifying the neural underpinnings of the inverted male/female LTTA phenotypes will be an important focus for future work.

Another key finding of this work is that LTTA in females is dynamically gated by the estrous cycle. Animals in the NR state show higher TZ avoidance, while SR animals display reduced avoidance, pointing to an interaction between reproductive drive and threat avoidance that carries an interesting ethological relevance: during periods of sexual receptivity, the drive to engage with the environment may appropriately shift the behavior away from avoidance. Our data identify progesterone, and not estrogen, as the key modulator of this effect: progesterone administration to ovariectomized females restores the reduced avoidance characteristic of the SR state, while estrogen has no detectable effect. Critically, this progesterone-dependent modulation requires melanopsin expression, as it is abolished in *Opn4^-/-^* animals. This indicates that progesterone may act by modulating ipRGC signaling directly, and indeed ipRGCs have been shown to express progesterone receptors. However, it is important to note that hormonal modulation is widespread and complex, and that sex hormones likely impact the LTTA circuit at multiple nodes.

Collectively, this work establishes the LTTA paradigm as a powerful tool for probing how internal states tune circuits that translate prior threat experience into long-term behavioral decisions. These findings show a previously unappreciated role for progesterone in modulating a retino-thalamic circuit to calibrate avoidance behavior and identify ipRGCs as a potential hormonal sensor within the LTTA circuit. More broadly, these findings highlight how the same circuit architecture can be differentially engaged across sexes to produce similar behavioral outcomes. This is an important consideration for understanding the neural basis of sex differences in threat-related behaviors and their relevance to conditions showing well-documented sex disparities in prevalence and symptomatology such as PTSD and anxiety disorders.

## Methods

### Animals

All procedures were approved by the Animal Care and Use Committee at Northwestern University (Protocol numbers: IS00003845 and IS00014627). Animals were housed in vivarium under 12:12 light/dark cycle conditions with *ad libitum* access to food and water. Temperature ranged from 21 to 23 °C and humidity ranged from 30% to 70%. P60-P120 female mice were used for this study. For behavior, chemogenetic and histological studies, we used *WT*, *Opn4^Cre^*(RRID:IMSR_JAX:035925)*, Opn4^Cre/+^; Vglut2^fl/fl^* (MGI:4879031) and *Gad2^Cre^* (IMSR_JAX:028867) female mice. For fiber photometry experiments we used WT female mice.

### Long-term threat anticipation paradigm

The Long-term threat anticipation (LTTA) paradigm was performed as described previously ^7^. The experimental arena consisted of a 40 x 40 x 40 cm^3^ opaque white foamboard Plexiglas box with a screen embedded in the ceiling. Throughout the experimental sessions, the monitor displayed a dim gray background (40 lux), on which a dark expanding disk was presented as the looming stimulus during the Exposure phase (Fig. 1A). We used Adobe Flash (RRID:SCR_017258) to design the looming stimuli and VLC Media Player (RRID:SCR_003070) to trigger the stimuli. Each stimulus consisted of ten expansions of a dark disk projected onto the gray background of the monitor. A single disc expansion consisted of two phases: 1) the disk expanded over a duration of 0.5 seconds, 2) the disk then remained static at its maximum expansion for 0.5 seconds before disappearing. An interval of 0.5-second was left before the next disk expansion; thus, each stimulus presentation lasted 10 seconds. Each mouse was exposed once, to one unique condition: 0, 6.6, 13.1, 27.5 or 36.5 % Michelson contrast. Michelson contrast was calculated based in arena illumination levels measured using a XL-500 BLE Spectroradiometer (nanoLambda Korea Co., Ltd).

Mice were dark-adapted for 20-30 min and then placed in the LTTA arena for 4 minutes (Pre-exposure, Fig. 1A). A single contrast of looming stimulus of contrasts ranging from 0-36.5% Michelson contrast and then recorded in the arena for 1 minute following the Exposure phase. After a retention period of 2 days in the home cage, 30-minutes dark-adapted animals were then placed in the same arena for 4 minutes with no stimulus triggered (Test phase, Fig. 1A) and behavior was recorded using two cameras (C920x HD, Logitech and ELP Wide Angle Infrared Webcam).

For chemogenetic experiments, 4 weeks after viral infections (see below), mice were dark adapted and 30 min before the Pre-exposure and Test phases of the LTTA paradigm, animals were i.p. injected with saline (Vehicle, 0.9 % NaCl) or Clonazepine N-Oxide (1 mg/kg) (CNO, Tocris Bioscience Cat# 6329). After i.p. injections behavioral assay was performed as described above. For local axon-terminal manipulations, 4 weeks after viral infections and cannula implantations (see below), mice were dark adapted and 20 min before the Pre-exposure and Test phases of the LTTA paradigm, animals were infused with saline (Vehicle, 0.9 % NaCl) or CNO (0.3 mg/kg). For Vehicle or CNO cannula infusions, bulk catheter (Cat#: BC-22, RWD Life Science Inc.) were attached to a Hamilton Syringe (Cat # 88500) to infuse a total volume of 0.3 μL at a rate of 0.1 μL/min. After infusions, behavioral assay was performed as described above. Mice with mistargeted viral expression or cannula implantation, confirmed by *post hoc* histology, were excluded from all analyses.

To study the role of sex hormones on LTTA, 4 weeks after ovariectomy (see below), mice were dark adapted and before the Pre-exposure and Test phases of the LTTA paradigm, animals were subcutaneously (s.c.) injected with Vehicle (Veh, corn oil) 1 hour previous the Pre-exposure phase, and Vehicle (Veh, corn oil), 17β-estradiol (3 μg/kg) (E2, Sigma-Aldrich E1024) or Progesterone (10 mg/kg) (P4, Sigma-Aldrich P0130) 1 hour previous the Test phase. After s.c. injections, the LTTA behavioral assay was performed as described above.

### Threat contrast sensitivity paradigm

The threat contrast sensitivity paradigm (TCS) was performed in the LTTA paradigm arena. As in LTTA, the monitor displayed a dim gray background (40 lux), on which a dark expanding disk was presented as the looming stimulus during the Exposure phase (Fig. 1A). Each stimulus consisted of three expansions of a dark disk projected onto the gray background of the monitor. A single disc expansion consisted of two phases: 1) the disk expanded over a duration of 0.5 seconds, 2) the disk then remained static at its maximum expansion for 0.5 seconds before disappearing. An interval of 0.5-second was left before the next disk expansion; thus, each stimulus presentation lasted 3 seconds. At least 30 s were left between consecutive stimulus presentations (Supplementary Fig. 4). Each mouse was exposed to three repetitions of: 6.6, 10.1, 13.1, 21.9, 27.5 or 36.5 % Michelson contras (18 looming stimuli total per animal). Michelson contrast was calculated based in arena illumination levels measured using a XL-500 BLE Spectroradiometer (nanoLambda Korea Co., Ltd). To trigger the looming stimuli, we used an adaptation of our custom Python-based algorithm ^9^.

### Behavior analysis

The LTTA behavior analysis was carried out as before^7^. Birefly, we used eZtrack (RRID:SCR_021496) automated tracking software ^21^ to analyze IR videos and quantify freezing behavior during Exposure phase of the LTTA paradigm at different looming stimuli contrasts. Freezing behavior was computed when mice were tracked immobile for at least 1 second. To quantify avoidance behavior, we used eZtrack software to analyze IR and regular light videos and quantify the time spent in the center of the arena during the Test phase (TZ_test_) of the LTTA paradigm. This parameter was normalized to the time spent in the center of the arena in the Pre-exposure phase for each mouse (TZ_pre-exp_).

To analyze TCS, we used DeepEthogram 22, a machine learning pipeline for supervised behavior classification during the Pre-exposure and Exposure phases of the TCS paradigm.

### Viral infection

Mice were anesthetized with isoflurane (Kent Scientific VetFlo system) and then unilaterally or bilaterally injected with 150 nl of AAVs (see Supplementary Table 1) in the perihabenular nucleus (PHb) (AP:-1.60, ML:±0.55, DV:2.65 mm) using a stereotaxic injector (Neurostar) controlled by the software Stereodrive at an injection rate of 30 nl/min. For intravitreal injections, mice were anesthetized by intraperitoneal (IP) injection of Avertin (2,2,2-Tribromoethanol) and a 30-gauge needle was used to open a hole in the *ora serrata*. Each eye was intravitreally injected with 1 μL of AAVs (see Supplementary Table 1) using a custom Hamilton syringe (Borghuis Instruments) with a 33-gauge needle (Hamilton). After surgery procedures, mice were subcutaneously injected with 20 mg/kg of meloxicam (Covetrus). Efficiency of the viral injections were tested *postmortem*.

### Estrous cycle tracking

Estrous cycle tracking in female mice was performed after the TCS paradigm and the Exposure and Test phases of the LTTA paradigm. Estrous cycle phases were determined using vaginal lavages and histological analysis of the smears as previously reported ^23^. Briefly, for vaginal lavage female mice were immobilized and ∼25-50 μl of PBS 1x was expelled at the opening of vaginal canal using a latex bulb attached with a filtered 200 μl tip. The fluid was then placed on a charged glass slide and the smears were dried at a hot plate. Smears were then stained with Cristal Violet 0.1% (Sigma, C0775) and imaged on a Leica SP5 microscope in bright field mode. Vaginal cytology was performed, and the relative ratio of cell types observed in smears were used to identify the stage of the estrous cycle of mice on the LTTA phase of sample collection. During Proestrus, cells are almost exclusively nucleated epithelial cells (Supplementary Figure 4). In Estrus phase, cells are predominantly cornified squamous epithelial cells (Supplementary Figure 4). During Metestrus, small darkly stained leukocytes predominate (Supplementary Figure 4) with fragments of cornified squamous epithelial cells. During Diestrus, predominating leukocytes (Supplementary Figure 4) and rare epithelial and cornified cells may still be present. For behavior analysis, mice were grouped into two categories: sexually receptive (SR - Proestrus and Estrus) and non-sexually receptive (NR - Metestrus and Diestrus) mice.

### Ovariectomy

Ovariectomy was performed as previously described ^24^. Briefly, mice were anesthetized with isoflurane (Kent Scientific VetFlo system). The site of the surgery was shaved and cleaned using 70% ethanol and povidone. Two small incisions were made on each flank. The fat pad was lifted outwards to exteriorize the ovary. The oviduct and the cranial part of the uterine horn distal to the ovary were crushed. The ovaries were removed and the oviduct was clamped for 15-20 seconds to minimize hemorrhage. The muscle wall was sutured using a 4-0 absorbable suture and the skin incision was closed using wound clips. During and 24 h after surgery procedures, mice were subcutaneously injected with 20 mg/kg of meloxicam (Covetrus). Sutures/wound clips were removed 7-14 days after surgery or when the incision was completely healed. Efficiency of ovariectomy procedure was tested by tracking the cessation of the estrous cycle as described above.

### Immunohistochemical procedures

For immunological studies in brain sections, mice were anesthetized by IP injection of Avertin and intracardially perfused with saline solution (PBS) followed by PFA 4% in PBS. Brians were dissected and post-fixated in PFA 4% in PBS overnight at 4 °C. For cryoprotection, brains were incubated in sucrose 30% for 3 days, embedded in OCT and then sectioned at 40 µm using a Leica CM1950 cryostat. For immunolabeling studies in retinas, animals were anesthetized by IP injection of Avertin and sacrificed by cervical dislocation. Eyecups were enucleated and retinas were dissected and then fixed in PFA 4% in PBS for 30-60 minutes at RT. Brain slices and retinas were washed and then blocked at 4 °C overnight in 6% normal goat serum in 0.3% Triton PBS prior to incubating in primary antibody solution for 2-3 nights at 4 °C. Then, tissue was washed and incubated in secondary antibody solution for 2 hours at RT (Supplementary Table 2). The tissue was finally mounted using Fluoromount (Sigma) and imaged using a Leica SP5 confocal microscope. Primary and secondary antibody solutions were made in 3% normal goat serum in 0.3% Triton PBS. For melanopsin and PHb-ipRGC mCherry expression analysis, high magnification images (pixel size of 0.36 µm) with a z-stack size of 1 µm were taken. For viral expression in brain regions, high magnification images (pixel size of 0.72 µm) with a z-stack size of 2 µm were taken. All images were processed and analyzed using ImageJ.

### Fiber photometry

Mice were anesthetized with isoflurane (Kent Scientific VetFlo system) and then unilaterally injected with 150 nl of AAV1-CAG-GCaMP6s-WPRE-SV40 (∼8 x 10^12^ GC/mL, see Supplementary Table 1) in the PHb using a stereotaxic injector (Neurostar) controlled by the software Stereodrive at an injection rate of 30 nl/min. A FP cannula (MFC_200/230-0.48_3mm_MF1.25_FLT, DORIC) was then implanted 150 µm above the site of injection. After surgery procedures, mice were subcutaneously injected with 2 mg/kg of meloxicam (Covetrus). Efficiency of the brain injection and fiber implantation were tested *postmortem*.

Behavioral assay and FP recordings were made 4 weeks post-surgery. For that, dark-adapted mice were exposed to the LTTA paradigm as described above. For FP recording, an optic fiber (MFP_400/430/1100-0.57_1m_FC-ZF1.25(F)_LAF, DORIC) was attached to the head of the animals and then were placed to the LTTA arena. FP signal was collected using an assisted electrical rotary system (AERJ_24_FMC, DORIC), Amplifier (Doric Fluorescence Detector, DORIC), LED driver (Laser Diode Fiber Light Source, DORIC) and processor (RZ5P Base Processor, Tucker-Davis Technologies). FP data was extracted using pMAT (RRID:SCR_022570) software ^25^. All FP data processing and behavior correlation analyses were made using eZtrack, Microsoft Excel (RRID:SCR_016137) and Graph Pad Prism 10.1.2 (RRID: SCR_002798).

### Statistical comparisons

All graphs and statistical analyses were performed using Graph Pad Prism 10.1.2. Normal distribution of data was tested using Shapiro-Wilk test. Data points were excluded only if identified as statistical outliers using the ROUT method (Q = 1%), which was pre-established prior to data analysis. No other data were excluded from the analyses. Two-sided Student’s t- or Mann Whitney U tests were used to compare two experimental groups. For multiple statistical comparisons, we performed one-way ANOVA followed by Tukey’s, Dunn’s, Kruskal-Wallis’ or Šídák’s tests. Significance was concluded when P < 0.05.

## Acknowledgments

National Institutes of Health grant R01 EY030565 and National Institutes of Health grant DP2 EY022584 (TMS). We thank Dr. Samer Hattar for the gift of *Opn4^Cre/+^* mice.

## Author contributions

Conceptualization: MLA and TMS. Methodology: MLA, CAB, SH, AES. Investigation: MLA, CAB, SH, AES and TMS. Visualization: MLA and TMS. Funding acquisition: TMS. Project administration: TMS. Supervision: MLA and TMS. Writing – original draft: MLA and TMS. Writing – review & editing: MLA and TMS.

---

**Supplementary Figure 1.**
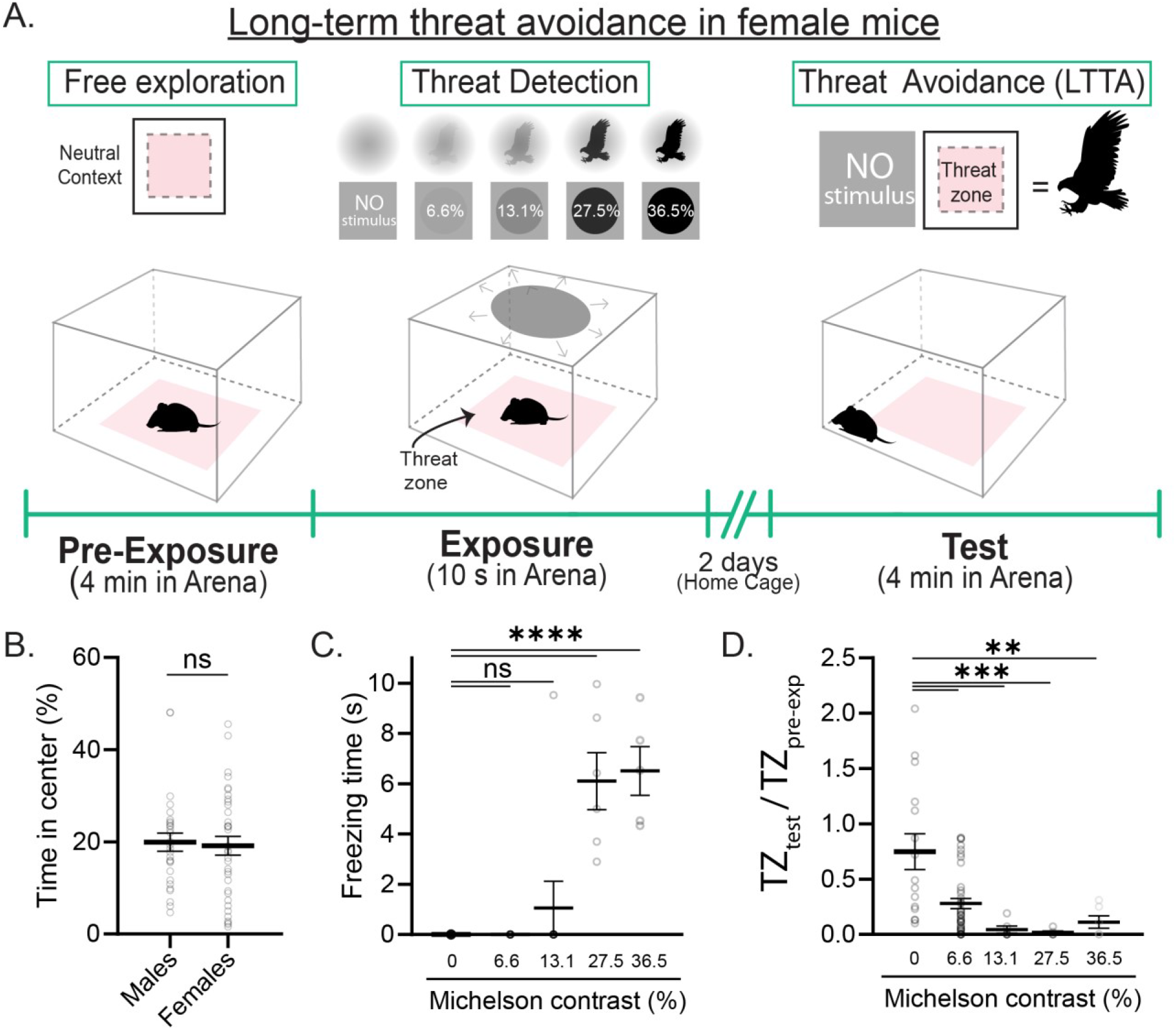
LTTA behavioral characterization of female mice. (A) Schematic representation of LTTA paradigm. (B) Percentage of time in center during the Pre-exposure phase in control male (n = 28) and female (n = 35) mice (P=0.476). (C-D) Freezing time during the Exposure phase (C) (0 vs. 6.6%, P=0.999; 0 vs. 13.1%, P=0.628; 0 vs. 27.5%, P=0.0001; 0 vs. 36.5%, P=0.0001) and time in threat zone (TZ) during the Test phase normalized to time in TZ during the Pre-exposure phase (D) (0 vs. 6.6%, P=0.0003; 0 vs. 13.1%, P=0.0007; 0 vs. 27.5%, P=0.0010; 0 vs. 36.5%, P=0.0023) at 0 (n = 15 mice), 6.6 (n = 38 mice), 13.1 (n = 6 mice), 27.5 (n = 5 mice) and 36.5% (n = 6 mice) Michelson contrast in control male mice. All data are Mean ± SEM, ns: P>0.05, ***P<0.001, ****P<0.001. Two-sided Student t-test and one-way ANOVA with Dunn’s multiple comparisons test.

**Supplementary Figure 2.**
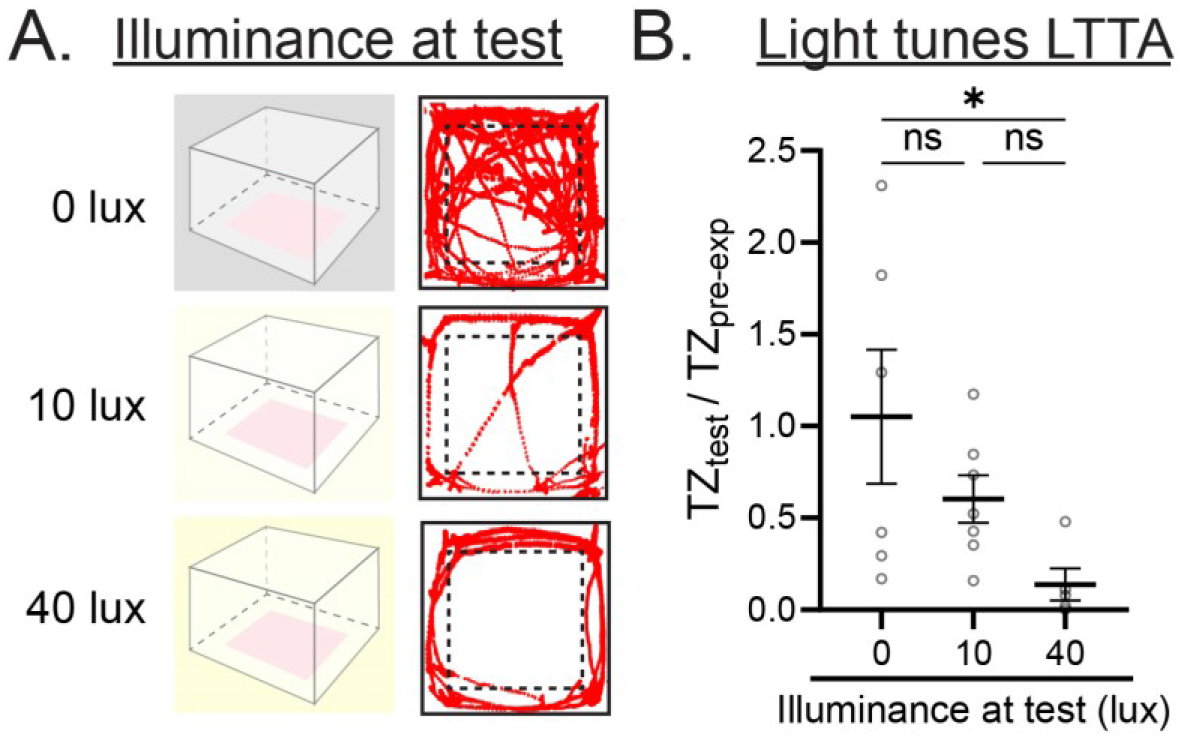
Female mice require visual input for LTTA. (A) Behavioral paradigm to test the role of light on LTTA (left) and representative animal position traces during the Test phase of the LTTA paradigm at 0, 10 and 40 lux of control female mice (right). (B) Time in TZ during the Test phase normalized to time in TZ during the Pre-exposure phase at 0 (n = 5 mice), 10 (n = 7 mice) and 40 (n = 6 mice) lux in control male mice (0 vs. 10 lux, P=0.369; 0 vs. 40 lux, P=0.045; 10 vs. 40 lux, P=0.356). All data are Mean ± SEM, ns: P>0.05, *P<0.05. One-way ANOVA with Dunn’s multiple comparisons test.

**Supplementary Figure 3.**
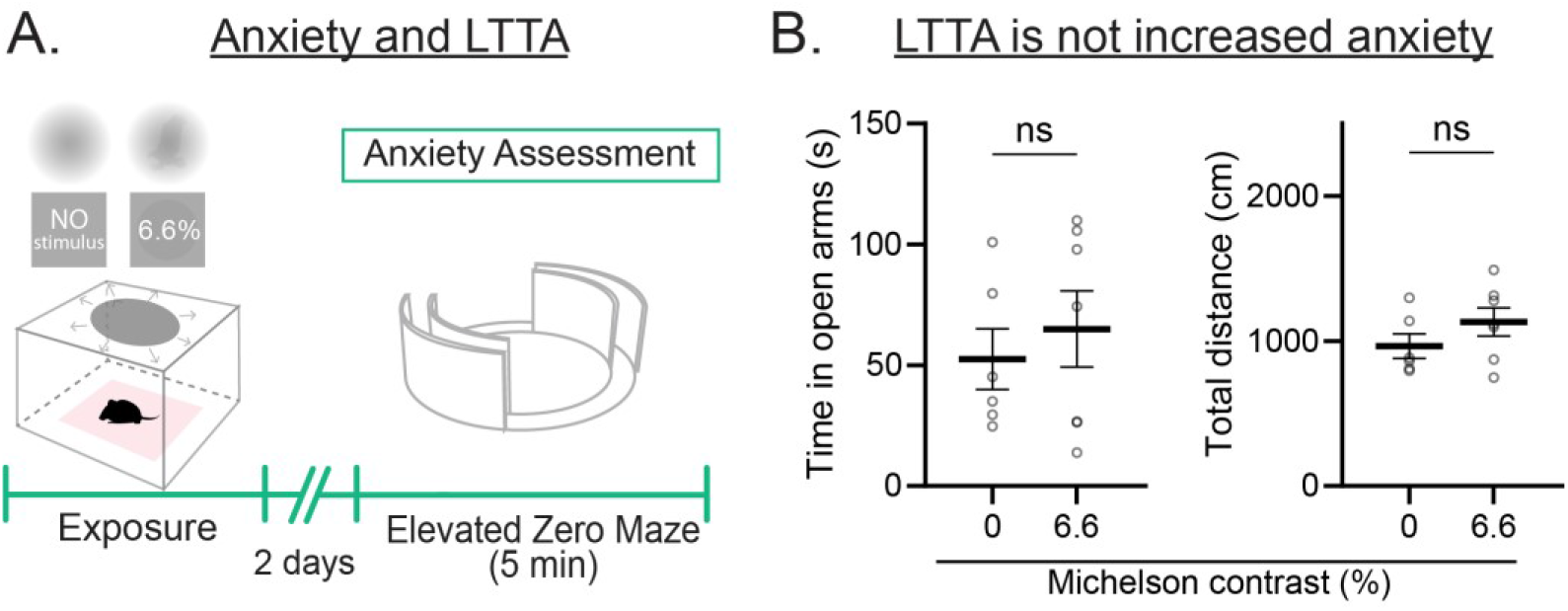
LTTA is not due to increased anxiety in female mice. (A) Behavioral paradigm to test anxiety in looming-exposed mice. (B) Time in open arms (left) (P=0.522) and total distance (right) (P=0.633) in the Elevated Zero Maze (EZM) assay of control male mice previously exposed to 0 (n = 6) or 6.6% (n = 7) looming stimulus. All data are Mean ± SEM, ns: P>0.05. Two-sided Student t-test.

**Supplementary Figure 4.**
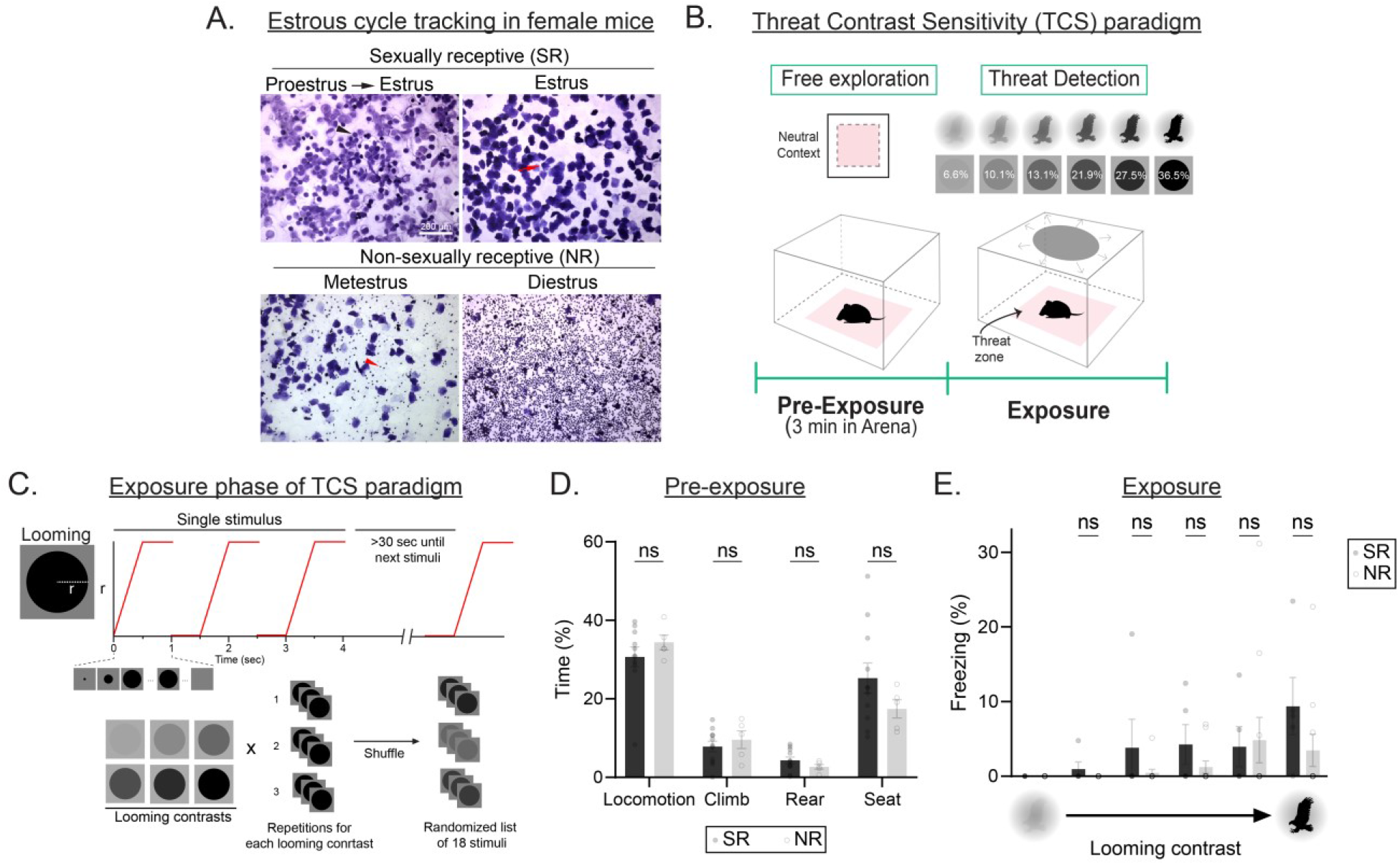
Estrous cycle does not impact looming stimuli detection. (A) Representative vaginal smears images of Sexually (SR) and Non-sexually (NR) receptive phases of estrous cycle of female mice. Black arrowhead: nucleated epithelial cell. Red arrow: cornified squamous epithelial cell. Red arrowhead: leukocyte. (B) Scheme of experimental timeline of the Threat Contrast Sensitivity paradigm (TCS). (C) Scheme of experimental timeline of looming stimuli display during the Exposure phase of the TCS paradigm. (D) Percentage of time of behavioral outcomes in the Pre-exposure phase of the TCS paradigm in control female mice in SR (n = 5) and NR (n = 11) stages of Estrus (Locomotion, P = 0.379; Climb, P = 0.491; Rear, P = 0.242; Seat, P = 0.214). (E) Freezing time during the Exposure phase of the TCS paradigm at 0, 6.6, 10.1, 13.1, 21.9, 27.5 and 36.5% Michelson contrast in SR (n = 5) and NR (n = 11) female mice (10.1%, P=0.939; 13.1%, P=0.966; 21.9%, P=0.910; 27.5%, P=0.999; 36.5%, P=0.781). All data are Mean ± SEM, ns: P>0.05. Two-sided Student t-test and two-way ANOVA with repeated measures with a Šídák’s multiple comparisons test.

**Supplementary Figure 5.**
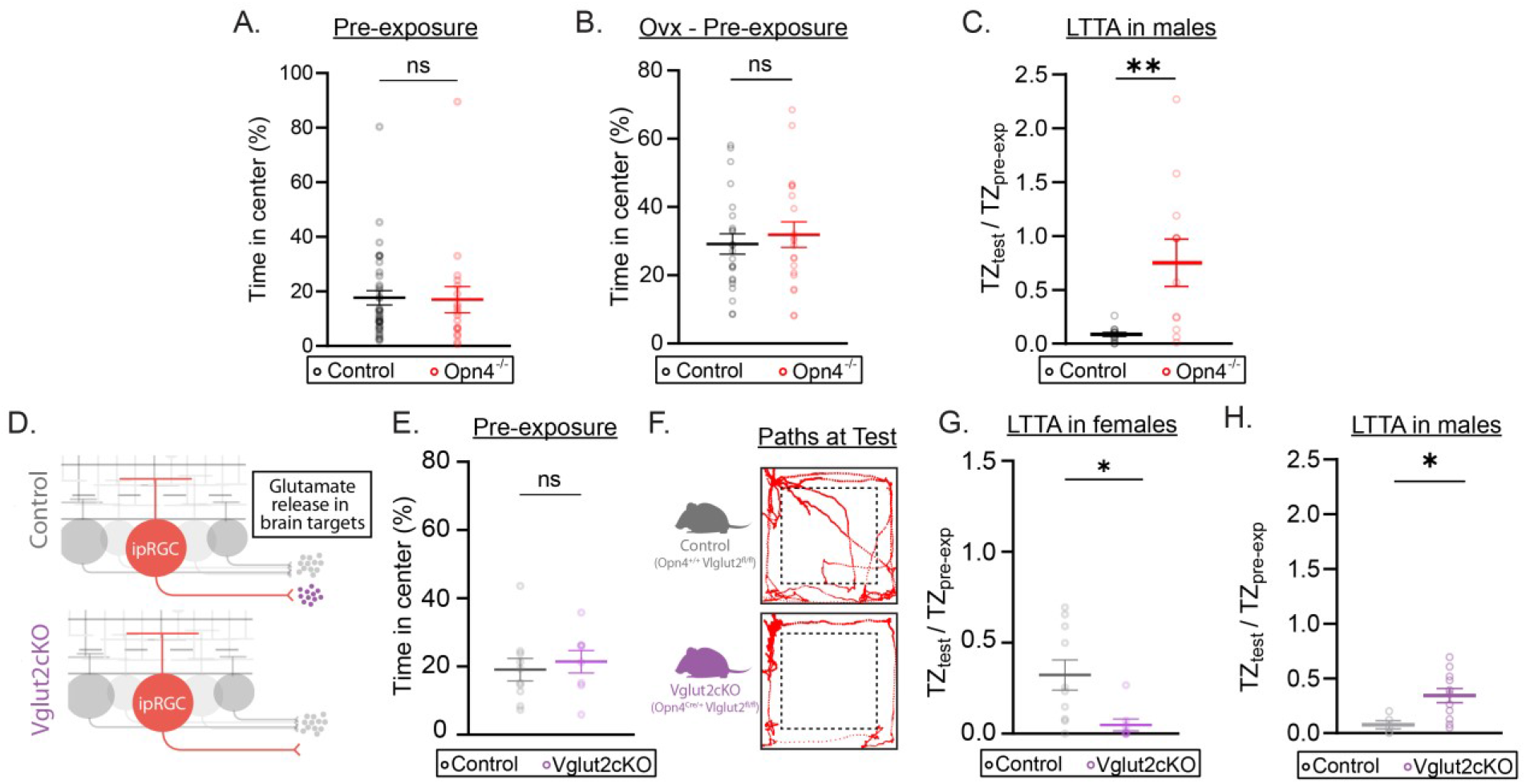
Ovariectomy does not impact baseline behavior in control and *Opn4^-/-^* mice. (A) Percentage of time in center during the Pre-exposure phase of control (n = 35) and melanopsin-null (*Opn4^-/-^*, n = 18) female mice (P=0.536) (B) Percentage of time in center during the Pre-exposure phase of ovariectomized (Ovx) (B) control (n = 23) and melanopsin-null (*Opn4^-/-^*, n = 20) female mice (P=0.506). (C) Time in threat zone (TZ) during the Test phase normalized to time in TZ during the Pre-exposure phase in *Opn4^-/-^*(n = 11, red) and control (n = 14, black) male mice exposed to 6.6% Michelson contrast (P=0.002). (D) Scheme of specific glutamate release ablation in ipRGCs. (E) Percentage of time in center during the Pre-exposure phase of Control (n = 10, grey) and Vglit2cKO (n = 8, purple) female mice (P=0.360). (F) Representative animal position traces of Vglut2cKO female mice during the Test phase of the LTTA previously exposed to 6.6% Michelson contrast. (G) Time in TZ during the Test phase normalized to time in TZ during the Pre-exposure phase in Vglut2cKO (n = 8, purple) and control (n = 10, grey) female mice exposed to 6.6% Michelson contrast (P=0.011). (H) Time in TZ during the Test phase normalized to time in TZ during the Pre-exposure phase in Vglut2cKO (n = 11, purple) and control (n = 5, gray) male mice exposed to 6.6% Michelson contrast (P=0.020). All data are Mean ± SEM, ns: P>0.05, *P<0.05, **P<0.01. Two-sided Student t- and Mann-Whitney U tests.

**Supplementary Figure 6.**
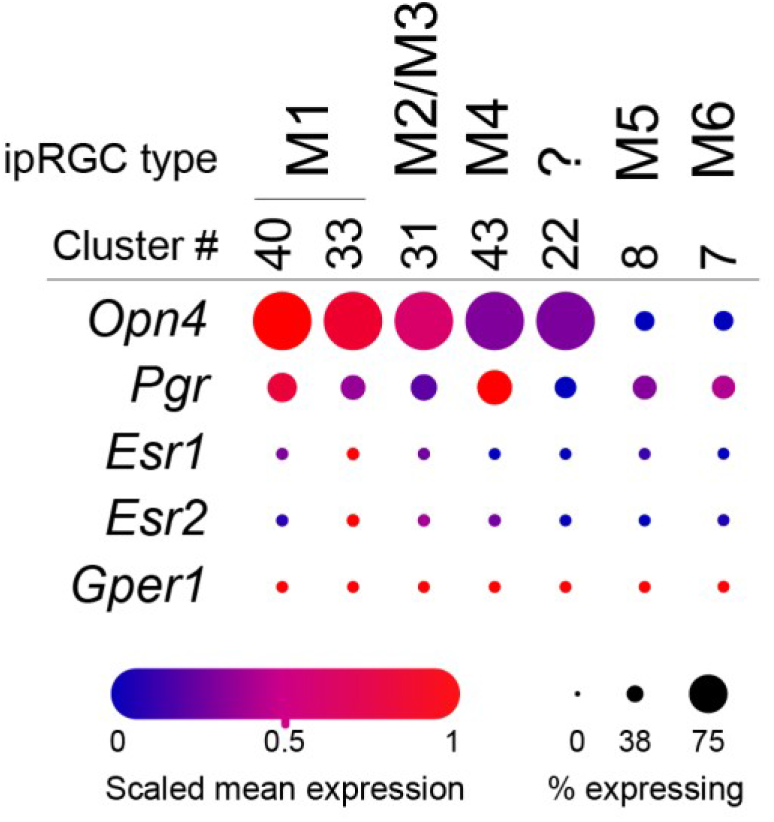
ipRGCs express progesterone, and not estrogen, receptors. mRNA levels of progesterone (*Pgr*) and estrogen (*Esr1*, *Esr2* and *Gper1*) receptor expression in ipRGC subtypes. Data from ^15^.

**Supplementary Figure 7.**
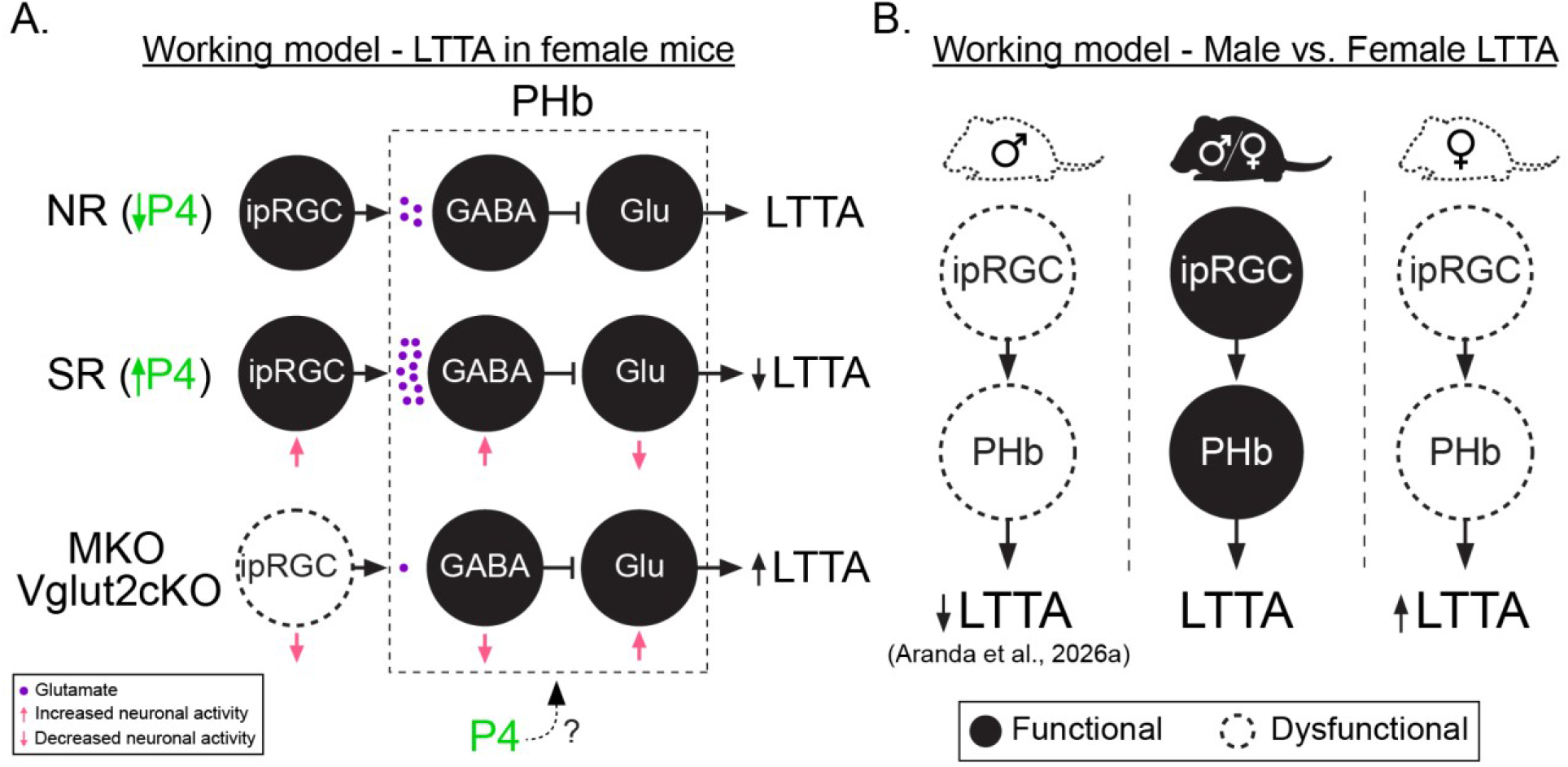
Working model for LTTA behavior in male and female mice. (A) Working model of progesterone in female LTTA. (B) Working model of LTTA circuit in male and female mice.

**Supplementary Table 1.**
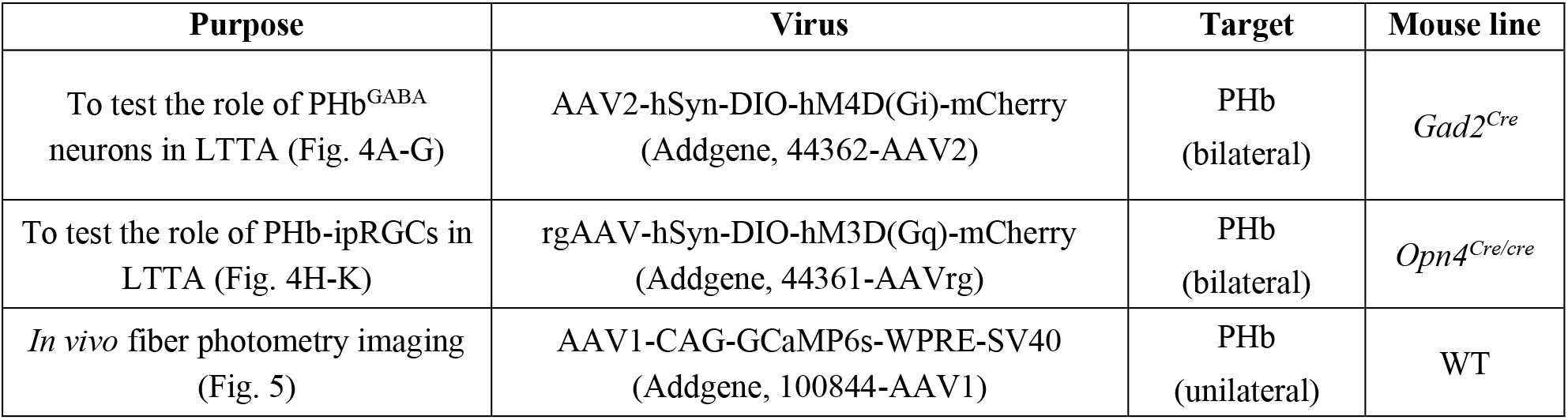
List of intersectional virus strategies used in this study.

**Supplementary Table 2.** List of antibodies used in this study. The dilution of all secondary antibodies was 1:500.

| Purpose | Primary Ab | Dilution | Secondary Ab |
| --- | --- | --- | --- |
| To characterize viral expression carrying mCherry (Fig. 4) | Rabbit dsRed (Takara, 632496) | 1:500 | Donkey anti-rabbit Alexa 594 (ThermoFisher, A-21207) |
| To characterize viral expression carrying GFP reporter (Fig. 5) | Chicken anti-GFP (Abcam, ab13970) | 1:500 | Donkey anti-chicken Alexa 488 (Jackson IRL, 703-545-155) |

